# Histone methyl-transferase G9a inhibition boosts the efficacy of immune checkpoint inhibitors in experimental hepatocellular carcinoma

**DOI:** 10.1101/2025.07.08.663670

**Authors:** Elena Adan-Villaescusa, Borja Castello-Uribe, Iker Uriarte, Eva Santamaria, Roberto Barbero, Miriam Belzunce, Amaya López-Pascual, Maria Ujue Latasa, Jasmin Elurbide, Emiliana Valbuena-Goiricelaya, Umberto Vespasiani-Gentilucci, Felipe Prosper, Antonio Pineda-Lucena, Bruno Sangro, Josepmaria Argemi, Pedro Berraondo, Pablo Sarobe, Maria Arechederra, Carmen Berasain, Alexis Cocozaki, Veronica Gibaja, Matias A Avila, Maite G Fernandez-Barrena

**Author notes:** Address for Correspondence*: Dr. Maite G Fernandez-Barrena. Hepatology Laboratory, Solid Tumors Program, CIMA. Pamplona, Spain. Both authors share senior authorship (Matias A Avila, Maite G Fernandez-Barrena).

## Abstract

**Background and Aims:** Immune checkpoint inhibitors (ICI) have revolutionized cancer therapy. Yet, their efficacy in hepatocellular carcinoma (HCC) remains limited, partly due to tumor-intrinsic mechanisms of immune evasion. This study focused on the identification of potential epigenetic drivers of immune resistance in HCC evaluating the therapeutic potential of targeting the histone methyltransferase G9a (EHMT2).

**Approach and Results:** We analyzed G9a expression across multiple human HCC cohorts and found that elevated G9a levels were inversely correlated with the most relevant immune-related gene expression signatures predictive of ICI responsiveness. Using HCC cell lines and orthotopic models implemented in immunocompetent mice, we assessed the effects of pharmacologic inhibition of G9a with two innovative epigenetic inhibitors, CM272 and EZM8266. G9a blockade enhanced tumor cell immunogenicity by restoring IFNγ responsiveness, increasing MHC-I surface expression, and promoting chemokine-mediated (CXCL10) recruitment of T cells. Mechanistically, G9a inhibition induced a viral mimicry response through derepressing endogenous retroviral elements and the accumulation of cytosolic double-stranded RNA. *In vivo*, G9a inhibition synergized with anti–PD-1 therapy to suppress tumor growth, significantly enhancing CD8⁺ T cell infiltration. Notably, in a clinically-relevant post-hepatectomy HCC recurrence model, the combination therapy overcame immune resistance.

**Conclusions:** G9a functions as a central epigenetic barrier to antitumor immunity in HCC. Pharmacologic G9a inhibition reprograms the tumor microenvironment, enhances immunogenicity, and sensitizes tumors to ICIs. These findings provide strong preclinical rationale for integrating G9a-targeted therapies with immunotherapy, particularly in perioperative settings.

## Introduction

Hepatocellular carcinoma (HCC), the predominant form of primary liver malignancy, ranks among the most prevalent and lethal cancers globally, with an incidence that continues to rise^1^. HCC typically arises in the context of chronic hepatic injury and sustained inflammation, most frequently due to hepatitis B or C viral infections, alcohol-associated liver disease, and increasingly due to metabolic dysfunction-associated steatotic liver disease (MASLD)^2^. Although various treatment modalities exist - including surgical resection, liver transplantation, and locoregional therapies - clinical outcomes remain suboptimal, particularly in patients presenting with advanced disease stages, for whom systemic therapy constitutes the primary therapeutic avenue. In recent years, immunotherapeutic strategies have markedly expanded the therapeutic landscape for HCC, particularly immune checkpoint inhibitors (ICI) targeting the PD-1/PD-L1 and CTLA-4 axes. Notably, the combination of atezolizumab (an anti–PD-L1 monoclonal antibody) with bevacizumab (an anti–VEGF-A antibody) has demonstrated superior efficacy over sorafenib in advanced HCC, establishing a new first-line standard with a median overall survival of 19 months^3^. Furthermore, ICI are actively being investigated in neoadjuvant and adjuvant contexts for early-stage disease, often in conjunction with surgery or local therapies^4^. Nevertheless, a substantial proportion of patients exhibit intrinsic resistance to ICIs, with only approximately 30% demonstrating objective clinical responses. Even among initial responders, many ultimately develop acquired resistance, limiting the durability of therapeutic benefit^5^. Thus, elucidating the mechanisms underlying immune evasion and identifying predictive biomarkers to guide patient stratification represent critical steps toward optimizing therapeutic efficacy in advanced HCC.

Various mechanisms have been implicated in both primary and acquired resistance, with increasing focus on the tumor microenvironment (TME), including immunosuppressive cell populations such as tumor-associated macrophages (TAMs), myeloid-derived suppressor cells (MDSCs), regulatory T cells (Tregs), inflammatory phenotypes, and T cell exhaustion^6^. Tumor cells further evade immune surveillance and cytolytic activity through complex genetic and epigenetic alterations^7^, which not only foster immune suppression but also interact with host-related factors such as diet, metabolism, and the gut microbiota^8^, ultimately contributing to immune resistance.

Epigenetic plasticity, including mechanisms such as DNA methylation, histone modification, and chromatin remodeling, plays a pivotal role in hepatocarcinogenesis, being strongly associated with tumor initiation, progression, and metastasis^9–12^. Importantly, emerging preclinical evidence suggests that specific epigenetic changes may modulate tumor–TME interactions and influence antitumor immune responses^13^. Although limited, emerging studies in murine HCC models have demonstrated that certain epigenetic modulators can affect tumor immune evasion and therapeutic resistance^14^. Consequently, pharmacological targeting of epigenetic regulators is being explored as a promising strategy to enhance the immunogenicity of HCC and improve ICI responsiveness^15^.

In this context, G9a (also known as EHMT2), a histone methyltransferase responsible for catalyzing mono- and dimethylation of histone H3 at lysine 9 (H3K9), is overexpressed in HCC and has been shown to orchestrate key oncogenic processes such as cell proliferation, survival, hypoxic adaptation, and metastasis^16,17^. Inhibition of G9a activity exerts potent antitumor effects in clinically relevant in vitro and *in vivo* HCC models, restoring tumor cell differentiation, but also mitigating the pro-oncogenic influence of the fibrotic stroma that dominates the HCC TME^16,18^.

In the present study, we demonstrate that elevated G9a expression in HCC patients is inversely correlated with transcriptional signatures previously associated with favorable responses to ICI-based therapies. We subsequently examined the efficacy of two selective G9a inhibitors, our tool compound CM272^16,18–20^ and the more clinically advanced molecule EZM8266^21^, in *in vitro* and *in vivo* HCC models, observing how G9a blockade reinstated several key steps of the cancer–immunity cycle, including the upregulation of endogenous retroviruses (ERVs) and the reactivation of immune signaling pathways and chemokines in tumoral cells that promote T cell infiltration. When combined with anti–PD-1 therapy, both inhibitors significantly suppressed tumor progression, in part by enhancing CD8+ T cell recruitment. These findings unveil a mechanistically grounded therapeutic combination capable of augmenting antitumor immunity and improving the efficacy of immunotherapy in HCC.

## Materials and Methods

### Transcriptomic data acquisition and preprocessing

Publicly available gene expression profiles of human HCC tissues were retrieved from their respective repositories. In the case of transcriptomic high-throughput (RNAseq) analysis, data corresponding to GSE114564^22^ and GSE148355^23^ were downloaded from the NCBI sequence data archive (SRA) in fastq format using SRA Toolkit Version 3.1.1. Adapter sequences and low-quality reads were removed using Trimgalore Version 0.6.0. with Qutadapt Version 1.18 ^24^. Reads were subsequently aligned to the hg38 reference genome using the splice-aware aligner STAR (version 2.7.9a)^25^. Gene level quantification was performed with STAŔs quantMode GeneCounts option to count the number of reads mapped to each gene. To ensure consistent and reliable gene expression data, MANE annotations (version 1.4)^26^ were utilized. TCGA-LIHC gene expression data generated by the TCGA research network (https://www.cancer.gov/tcga) was retrieved as STAR-counts aligned to the hg38 genome, using TCGAbiolinks R package (version 2.34) in R software Version 4.4.2 (hereafter called R)^27^. Microarray data for GSE14520-GPL3921^28^, GSE89377^22^ were directly downloaded from the processed matrices available in the NCBI repository, for which the gene symbols for each platform were updated using AnnotationDbi version 1.68.0 and org.Hs.eg.db version 3.20.0 packages. For all RNA-seq datasets, including TCGA-LIHC, raw counts were normalized using the trimmed mean of M-values (TMM) using edgeR (version 4.4.1). TMM normalization accounts for library size differences and composition biases, ensuring accurate comparisons between samples.

### Molecular classification and gene expression analysis

To evaluate transcriptomic differences, first, low expression genes were filtered using the filterByExpr function implemented in edgeR, with the default setting. After that, the voom-limma pipeline for differential expression was used. For the assesement of the enrichment of specific genesets, single sample geneset enrichment analysis algoritm (ssGSEA) from the corto package (version 1.2.4)^29^ was used. Stratification of the patients into high and low subgroups for each of the signatures has been done by using the median of the ssGSEA score. Gene Set Enrichment Analysis (GSEA) was performed in pre-ranked mode using the clusterProfiler R package, with genes ranked by log2 fold change. Gene sets were sourced from the Hallmark and Gene Ontology (GO) collections of the Molecular Signatures Database (MSigDB) via the msigdbr package.

### Cell culture, treatments and reagents

The murine HCC cell line NM53 was derived from tumors generated in mice by hydrodynamic tail vein injections of a transposon vector expressing MYC (pT3-EF1a-MYC), a vector expressing SB13 transposase (CMV-SB13) and a CRISPR-CAS9 vector expressing a single-guide RNA (sgRNA) targeting p53 (px330-sgp53)^30^. PM299L murine HCC cell line, kindly provided by Dr. A. Lujambio (Icahn School of Medicine, Mount Sinai, NY, USA) was also derived from tumors generated in mice by hydrodynamic tail vein injections of the pT3-EF1a-MYC) MYC expressing vector, the CMV-SB13 vector and a transposon vector expressing activated β-catenin (CTNNB1-N90), which presents a deletion of the 90 first amino acids leading to constitutive activation^30^. NM53, PM299L, and human HuH7 and PLC/PRF5 cell lines were cultured as described^16^. Treatment times and dosages in the different experiments are specified throughout the manuscript, with controls receiving equivalent concentrations of dimethyl sulfoxide (DMSO) (always < 0.1% of the final volume). All cells were routinely tested for mycoplasma.

CM272 was synthesized as described previously^31^. EZM8266 was provided by Epizyme (an Ipsen company). The structure and synthesis of EZM8266 are described in the patent as “Compound 5R” (Campbell JE, Duncan KW, Mills JEJ, Munchhof MJ. Amine-substituted heterocyclic compounds as Ehmt2 inhibitors, salts thereof, and methods of synthesis thereof. 2019. Available from: https://patentscope.wipo.int/search/en/detail.jsf?docId¼WO2019079540&_cid¼P10-LS1T2D-13876-1.). Recombinant human and mouse IFNγ was from R&D Systems (Minneaopolis, MN, USA). The Anti-PD1 antibody (clone RMPI-14) and the isotype control antibodies are from Bio X Cell (Lebanon, NH, USA).

### RNA isolation, quantitative real-time PCR (RT-qPCR) and RNA sequencing (RNAseq)

Total RNA was isolated from the cell lines using the automated Maxwell system (Promega, Madison, WI, USA). Quantitative reverse transcription PCR (qRT-PCR) was carried out as previously described, and gene expression levels were normalized to the housekeeping gene H3F3A^16,20^. Primer sequences are available upon request.

RNA concentration and integrity were assessed using the Qubit High Sensitivity RNA Assay Kit (Thermo Fisher Scientific, Waltham, MA, USA) and the 4200 Tapestation system equipped with High Sensitivity RNA ScreenTape (Agilent Technologies, Santa Clara, CA, USA). All RNA samples exhibited high integrity, with RNA Integrity Number (RIN) values exceeding 8. Library construction was carried out using the Illumina Stranded mRNA Prep Ligation Kit, following the manufacturer’s instructions (Illumina, San Diego, CA, USA). For each sample, 100 ng of total RNA was used. RNA sequencing was conducted as previously described^19^ at the Genomics Unit of the Center for Applied Medical Research (CIMA), University of Navarra, Pamplona, Spain.

### RNAseq Analysis

RNAseq data from both human and mouse samples have been first preprocessed using the standardized workflow described previously. For human HCC cell lines PLC/PRF/5 reads were aligned to the hg38 reference genome using STAR and the MANE Annotations. In the case of mouse NM53 cell line experiments, reads were aligned to the GRCm39 genome using STAR and the GENCODE vM37 GTF annotation. The resulting raw counts were normalized using the trimmed mean of M-values (TMM) using edgeR (version 4.4.1). Differential expression analysis and all subsequent downstream analyses, including pre-ranked Gene Set Enrichment Analysis (GSEA), were conducted according to the procedures described above.

### Identification and differential analysis of Transposable Elements (TE)

To quantify transposable element (TE) expression at the family level, RNA-seq samples were first trimmed using TrimGalore, then aligned to the mouse genome (GRCm39) with STAR, using the corresponding GENCODE vM37 GTF annotation. The resulting unsorted BAM files were used as input for TEtranscripts, which was run in multimapping mode with both the GENCODE vM37 GTF and a pre-generated TE annotation file (GRCm39 GENCODE rmsk TE GTF) to assign read counts to genes and TEs. The generated count tables were then preprocessed by filtering out low-expression features and normalized using the TMM method. Differential expression analysis was performed using edgeR.

### Colony formation, anchorage-independent growth, migration and invasion assays

Colony formation assays were conducted using the specified cell lines following previously established protocols^16^. A total of 3,000 cells were plated in six-well plates containing complete medium, and treated with EZM8266 (at indicated doses) the following day. Media were refreshed every two days, and cultures were maintained for approximately 15 days until observable differences emerged between treatment conditions. Cells were then washed with PBS, fixed with 4% formaldehyde (Sigma-Aldrich) in PBS for 10 minutes, and stained using crystal violet. Representative images were captured. Each assay was carried out in at least two independent biological replicates, each with three technical replicates.

The soft agar assay was performed to evaluate anchorage-independent cell growth. Briefly, a base layer of 0.6% agar in complete culture medium was prepared in six-well plates. Once solidified, 5,000 PLC/PRF5 cells per well were suspended in 0.3% agar mixed with complete medium and plated on top of the base layer. Plates were incubated at 37 °C in a humidified CO₂ incubator for 3 weeks in the presence or absence of EZM8266. At the endpoint, colonies were stained with crystal violet and counted under the microscope (Leica, Wetzlar, Germany). Experiments were conducted in triplicate For migration and invasion assays, HuH7 cells (10⁵) were seeded into the upper chambers of Transwell inserts (Corning, Glendale, AZ, USA) featuring 8.0 μm pore polycarbonate membranes^19^. In both assays, cells were cultured in medium containing 0.5% FBS, while the lower chambers were filled with medium containing 30% FBS to serve as a chemoattractant. Following cell attachment, EZM8266 was applied at the indicated concentrations for 24 hours. Membranes were then washed with PBS, fixed with 4% paraformaldehyde for 24 hours, rinsed again with PBS, and stained with 0.5% crystal violet in 2% methanol. Each experiment was performed in triplicate. Between 3 and 5 images per Transwell insert were acquired at 5X magnification using an inverted microscope (Leica, Wetzlar, Germany). ImageJ software^32^ was used for image analysis and quantification.

### Enzyme-linked immunosorbent assay (ELISA)

CXCL9 and CXCL10 concentrations were measured in culture supernatants collected at the end of the incubation periods using commercial ELISA kits for mouse (CXCL9: R&D Systems, DY492; CXCL10: R&D Systems, DY466) and human cell lines (CXCL10: BD Biosciences, 550926). Assays were performed according to the manufacturers’ instructions.

### Flow cytometry

After treatments cells detached from culture plates by gentle trypsin treatment, cells were pelleted and washed with phosphate-buffered saline (PBS). Dead cells were stained with Zombie NIR™ viability dye (BioLegend, 423105). To block Fc receptors, cells were incubated with purified anti-mouse CD16/32 antibody (clone 93, BioLegend, 101302). Surface staining was performed using anti-mouse H-2K^b^ antibody (clone AF6-88.5, BioLegend, 116505). Finally, cells were fixed with BD Cytofix™ Fixation Buffer (BD Biosciences). Flow cytometry was conducted using a CytoFLEX flow cytometer (Beckman Coulter), and data were analyzed with FlowJo software (version 10.8).

### Immunofluorescence

For immunofluorescent detection of dsRNA cells were cultured on coverslips. After overnight incubation, cells were treated with CM272 or EZM8266 at the indicated doses. Immunofluorescence was performed as previously reported ^33^. Briefly, after 48h treatment, cells were fixed with ice-cold methanol for 15 min at room temperature and washed twice with PBS. Cells were permeabilized with 0,2% Triton X-100 for 10 min at RT. After washing, coverslips were blocked with Superblocking buffer (Thermo Fisher Scientific) for 1 h at room temperature and incubated 1h at RT with anti-dsRNA antibody 9D5, ab00458-2.3 (Absolute Antibody, Cleveland, UK) diluted in 1% BSA in PBS. After washing with 1% BSA in PBS, cells were incubated with fluorophore-conjugated secondary antibody in 1% BSA in PBS for 1 h at room temperature, washed and stained with vectashield (Vector laboratories, Burlingame, CA, USA) containing DAPI. Images were obtained using the Zeiss Axio Imager.M1 microscope (Zeiss, Oberkochen, Germany).

### *In vivo* experiments

All animals received humane care according to the ’Guide for the Care and Use of Laboratory Animals’ written by the National Academy of Sciences and published by the National Institutes of Health (NIH publication 86-23, revised 1985). Protocols were approved by the Animal Care Committee of the University of Navarra and were performed following their guidelines (ethical committee approval **# 048/22**). Animal experiments followed the Animal Research: Reporting of *In Vivo* Experiments (ARRIVE) guidelines (http://www.nc3rs.org.uk/arrive-guidelines), developed by the National Centre for the Replacement, Refinement and Reduction of Animals in Research (NC3Rs) to improve standards and reporting of animal research. For all studies 6- to 8-week-old male C57BL/6J mice purchased from Jackson Laboratories (Bar Harbor, ME, USA) were used. For the orthotopic tumor model, subcutaneous tumors were first generated with PM299L cells in C57BL/6J mice as described previously^16^. When tumors reached approximately 1 cm in diameter, animals were sacrificed and tumor tissues were sliced into equal fragments of ∼1 mm³. These fragments were orthotopically implanted into the left liver lobes of two groups of C57BL/6J mice via laparotomy. Tumor engraftment was monitored by ultrasound scan (US) using the Vevo 770 High-Resolution Imaging System (VisualSonics, Toronto, Canada), enabling *in vivo* visualization, assessment, and measurement of tumors. When lesions reached ∼2 mm³, mice (n = 5 per group) were randomized into control and treatment groups. Mice received 300 mg/kg (oral gavage) of EZM8266 or the same volume of vehicle (0.1% Tween80 and 0.5% methylcellulose, both from Sigma-Aldrich, St. Louis, MO, USA, in sterile water) for the indicated period of time. Mice were then sacrificed and tumors extracted. In a second orthotopic HCC model, 25.000 PM299L cells were intrahepatically injected in the left liver lobe of C57BL/6J mice. At the indicated times, mice were randomly divided into 4 groups and treated with vehicle (0.5% Methylcellulose+0.1% Tween-80 and human igG isotype, Bio X Cell), EZM8266 (300 mg/kg), anti-PD-1 (RMP1-14, Bio X Cell, 50 µg per mouse), or EZM8266 plus anti-PD-1 for 4 weeks. EZM8266 was administrated via oral gavage 5 days per week. Anti-PD-1 was administrated via intraperitoneal injection twice per week. Then, mice were sacrificed and liver tumors extracted. Finally, we implemented a third orthotopic syngeneic experimental model in which the growth of implanted HCC cells is stimulated by a concomitant partial hepatectomy (PH). This model mimics the frequent early tumor recurrence observed in patients undergoing HCC resection^34–36^ . To this end, 5.0 × 10^4^ PM299L cells in 0.2 mL of PBS were directly injected into the inferior right hepatic lobe remnant after a 33% PH performed essentially as previously described^37^. At the indicated times, mice were randomly divided into 4 groups and treated with vehicle control, CM272 (5 mg/kg), anti-PD-1 (RMP1-14, Bio X Cell, 50 µg per mouse), or CM272 plus anti-PD-1 for 4 weeks. CM272 was administrated via intraperitoneal injection 5 days per week. Anti-PD-1 was administrated via intraperitoneal injection twice per week. At the indicated time point mice were sacrificed and liver tumors extracted.

### Biochemical parameters

Serum levels of alanine aminotransferase (ALT), aspartate aminotransferase (AST), and lactate dehydrogenase (LDH) were measured using a C311 Cobas Analyzer (Roche Diagnostics GmbH, Mannheim, Germany) following the manufacturer’s instructions.

### Immunohistochemical analyses

Immunohistochemical analyses on liver and HCC tissues were performed essentially as described ^16,18,19^. Antibodies used were: 98941T from Cell Signaling (Danvers, MA, USA) for CD8, and ab183685 from Abcam (Cambridge, UK) for CD4, both at 1:100 dilution. HRP conjugated Envision secondary antibody K4003, followed by DAB reagent K3468, DAKO, both from DAKO (Glostrup, Denmark), were applied for detection. For signal quantification, images were analyzed with the QuPath software v0.3.217. Tissue sections were counterstained with Hematoxylin (Sigma-Aldrich) and dehydrated. Negative controls were performed omitting primary antibodies.

### Statistical Analyses

Genes were considered differentially expressed if the adjusted p-value, calculated using the Benjamini-Hochberg FDR method, was below 0.05. For correlation analysis of gene expression data Spearmańs correlation was used. Statistical analyses were performed using GraphPad Prism software (v9; GraphPad Software Inc., La Jolla, CA, USA). For comparison between two groups, paired two-tailed Student’s t-test, or Kruskal-Wallis ANOVA-test were used according to sample distribution. All reported p values were two-tailed and differences were considered significant when p<0.05.

## Results

### G9a expression negatively correlates with prognostic molecular signatures of ICI response in HCC patients

Several transcriptomic classification schemes have been developed to delineate HCC subtypes according to their biological behavior, molecular characteristics, and clinical manifestations. These systems incorporate key parameters such as proliferative activity, oncogenic pathway activation, differentiation status, mutational landscape, and immune microenvironmental features^38,39^. In particular, immune-related transcriptomic profiling has recently identified discrete HCC immune phenotypes—namely, immune-inflamed, immune-excluded, and immune-desert—each associated with distinct responses to ICI-based therapies and with differing prognostic implications^40^. Several gene expression signatures reflecting inflammatory signaling or inflamed HCC states have been shown to correlate with favorable responses to anti-PD-1 monotherapy^41–43^. Consistently, biomarkers of pre-existing antitumor immunity—such as high PD-L1 expression, enrichment of T-effector cell signatures, and elevated intratumoral CD8⁺ T cell densities—have been associated with improved clinical outcomes in patients receiving the combination of atezolizumab and bevacizumab. From these findings, a composite “atezo+bev response signature” (ABRS) was derived to stratify responders within this therapeutic setting^44–46^.

To explore whether expression of the histone methyltransferase G9a may influence response to ICIs in HCC, we first evaluated G9a transcript levels across The Cancer Genome Atlas (TCGA) HCC dataset and four additional independent HCC cohorts. Across all datasets analyzed, G9a expression demonstrated a consistent and statistically significant negative correlation with multiple gene signatures predictive of ICI response, including the Inflamed ^43^, IFNAP ^41^, IFN18^47^ and the mentioned ABRS signatures^46^. When HCC patients are stratified according to the score of each predictive gene signature, those with lower signature scores—indicative of a poorer likelihood of response to immunotherapy—consistently exhibit higher levels of G9a expression **(Fig. 1A** and **Suppl Fig.1A)**. Interestingly, a recent study employing single-cell RNA sequencing (scRNA-seq) of advanced HCC specimens identified 21 distinct cell-type-specific gene signatures. The potential utility of these signatures as predictive biomarkers of response to atezo+bev therapy was evaluated using bulk transcriptomic data from pretreatment advanced HCC samples. Different molecular subtypes of responders were delineated, one subgroup being characterized by the combined intraumoural presence of two CD8+ effector T-cell subtypes and CXCL10+macrophages, representing an immune-rich TME^45^. Notably, when applying these refined response-specific signatures across the same independent cohorts, G9a expression again showed a significant inverse correlation, further supporting the notion that patients with higher G9a expression levels consistently exhibit lower predictive signature scores **(Fig. 1B** and **Suppl Fig.1B)**. We also confirmed, both in the TCGA dataset and across other four independent HCC cohorts analyzed, how G9a expression exhibits in most of the cases a statistically significant negative correlation with multiple individual genes comprising each of the evaluated predictive signatures **(Fig. 1C** and **Suppl Fig. 1C)**. Collectively, these data implicate G9a as a potential negative regulator of antitumor immune responses in HCC and suggest its involvement in limiting patient responsiveness to ICI-based therapies.

**Figure 1.**
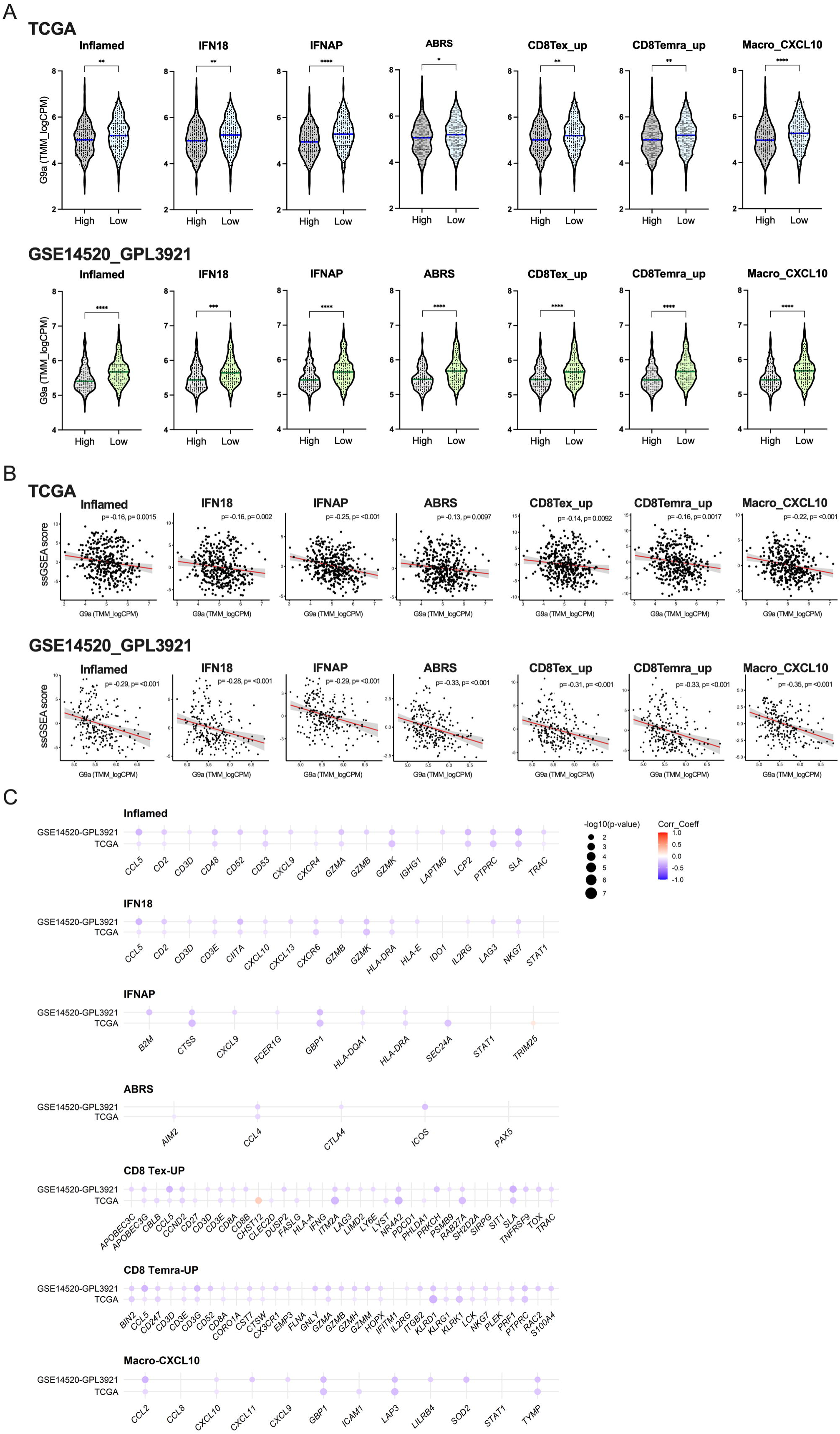
G9a expression in HCC tissues is associated with gene signatures predictive of immune-based therapy response. (A) G9a expression in patients from the TCGA and GSE14520_GLP3921 HCC gene expression datasets stratified according to the score of the indicated predictive signatures. (B) Spearman correlation plots showing the inverse association between G9a gene expression and the indicated signatures, as quantified by ssGSEA scores. Correlation coefficients (Spearman’s ρ) and corresponding p-values are indicated in each plot. (C) Correlation between G9a expression and that of the different genes that constitute the indicated gene signatures in the TCGA and GSE14520_GLP3921 HCC gene expression datasets. ** p<0.01, *** p<0.001, **** p<0.0001.

### G9a inhibition enhances the immunogenicity of HCC cells

Among both tumor-intrinsic and tumor-extrinsic mechanisms implicated in resistance to immunotherapy, deficiency in interferon-gamma (IFNγ) signaling has emerged as a critical contributor to immune evasion^7^. This impairment disrupts several key steps of the antitumor immune response, including effector immune cell trafficking and infiltration, antigen presentation, and tumor cell recognition^48^. In HCC, IFNγ signaling output is notably reduced compared to other solid malignancies^49^. Importantly, elevated baseline expression of IFNγ-responsive genes (IRGs) has been associated with favorable clinical responses to ICI therapies^41^. To explore the tumor-intrinsic regulatory mechanism of G9a on immunity, RNA-seq analyses were performed in NM53 cells, a murine HCC cell line, treated with our selective epigenetic inhibitor CM272 in the absence or presence of IFNγ. We observed multiple changes in gene expression in NM53 cells in both conditions **(Fig. 2A).** Consistently, differentially expressed genes were involved in biological functions previously associated with the inhibition of G9a activity in HCC, such as oxidative phosphorylation and mitochondrial activity^16^. In addition, we identified genes involved in numerous biological processes related to immune pathways, including responses to cytokines or to interferon **(Fig. 2B** and **Suppl Fig. 2A)**. Gene set enrichment analysis (GSEA) of differentially expressed genes highlighted various functional categories such as IFNγ response (*Stat1, Cxcl10*), innate immune response (*Tap1, Psme1*), response to virus (*Irgm2, Cxcr4, Cgas*) and antigen processing and presentation (*B2m, Tap1, Psme1*) signaling pathways that were significantly enriched in CM272-treated cells. Interestingly, all these pathways, were overrepresented in NM53 cells concomitantly stimulated with IFNγ and CM272 **(Fig. 2C)**. To validate the above results, a second murine HCC cell line, PM299L^30^, was treated with CM272 in the presence or absence of IFNγ. RT-qPCR analysis of the expression of key selected genes confirmed the effects of G9s inhibition on the response of HCC cells to INFγ (**Fig. 2D**). Importantly, similar results were found in human HCC cells **(Suppl Fig. 2B)**. In summary, these results support the notion that G9a is involved in the repression of IFNγ-responsive genes in human and mouse HCC cells.

**Figure 2.**
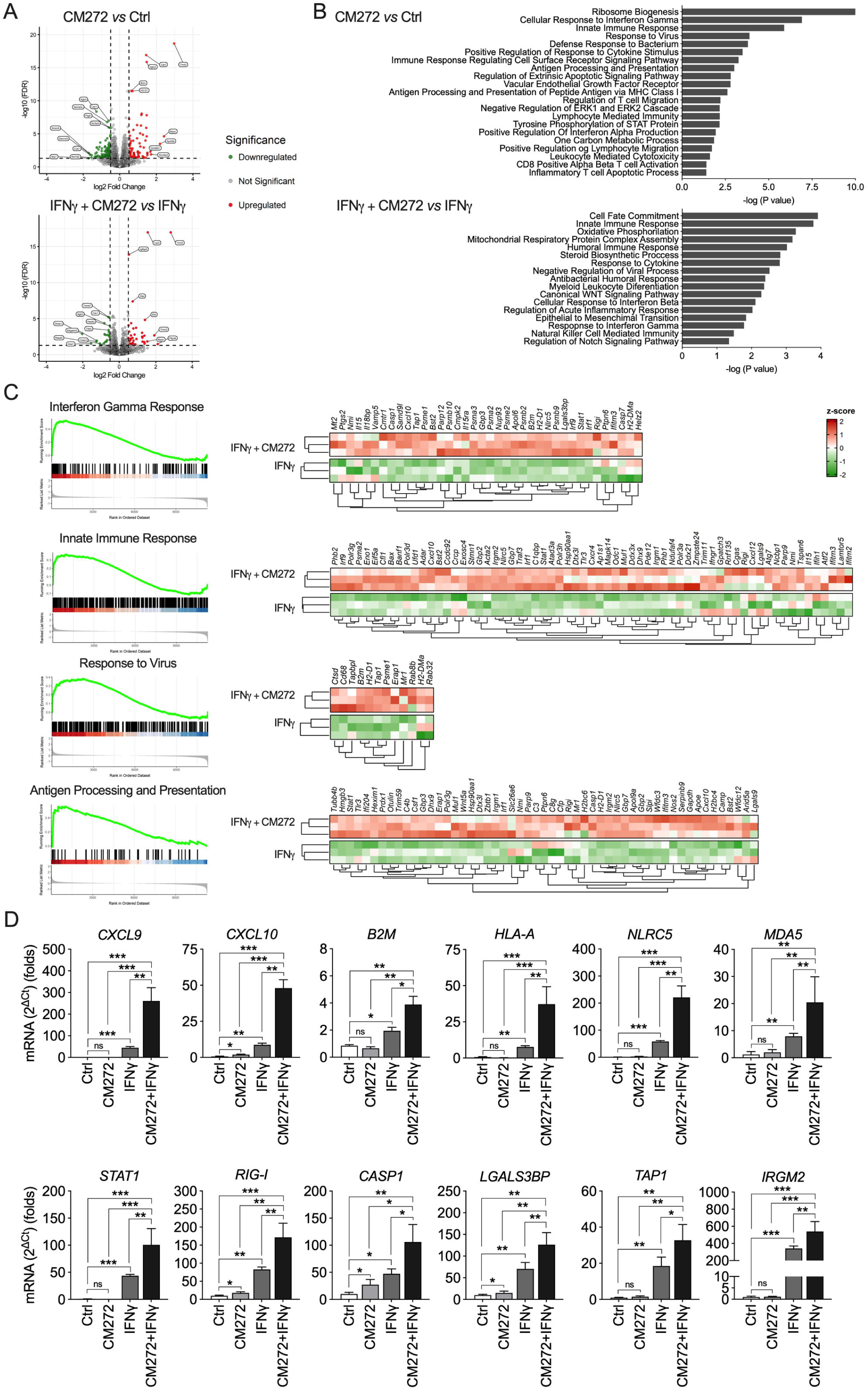
G9a inhibition enhances the immunogenicity of HCC cells. (A) Volcano plots representing all the genes with significant differential expression (p<0.05) in NM53 murine HCC cells treated with CM272 (400 nM for 48 h, upper panel) *vs* control (Ctrl) cells, or with CM272 (400 nM) for 24 h and then with INFγ (75 U/mL) for another 24 h *vs* INFγ alone (75 U/mL for 48 h, lower panel), as analyzed by RNA-seq. (B) Most relevant categories of differentially expressed genes identified by Gene Ontology Biological Process (GO-BP) functional classification in NM53 cells treated as indicated before. (C) GSEA analysis of specific categories including heatmaps with a list of genes modulated by INFγ *vs* INFγ plus CM272 selected from the RNA-seq data. (D) Validation of the effects of CM272, INFγ and their combination, as indicated above, on the expression of selected genes identified in the RNA-seq analyses in the murine HCC cell line PM299L. * p<0.05, ** p<0.01, *** p<0.001.

## Pharmacological inhibition of G9a with EZM8266 displays antitumoral properties in HCC

Currently, there are limited options for a safe and efficacious *in vivo* targeting of G9a. However, EZM8266 was developed by Epizyme-Ipsen as an orally available G9a inhibitor with promising pharmacodynamic and pharmacokinetic properties^21^. To evaluate the potential antitumor properties of this new compound in the context of HCC, we conducted colony formation, anchorage-independent growth and transwell assays in different human HCC cell lines. EZM8266 treatment markedly decreased colony formation, cell migration and cell invasion in all cell lines tested (**Fig. 3A-D**). To gain insight into the mechanisms of the antitumoral effects of EZM8266, we performed RNAseq analyses in a human HCC cell line. We detected 1069 upregulated and 1120 downregulated genes compared with controls (P < 0.01). Gene ontology (GO) functional classification of differentially expressed genes identified general categories linked to the regulation of diverse functions already associated with G9a activity, such as double strain DNA damage repair, autophagy or L-serine metabolic processes ^16,50–53^. Similarly, as observed in mouse HCC cells, G9a inhibition significantly and substantially increased the expression of many genes involved in innate immune response, interferon response, or processes such as dsRNA processing related to immunity. **(Fig. 3F, G)**. We next treated the murine HCC cells PM299L with EZM8266 in the absence or presence of IFNγ. We observed virtually the same changes as previously seen with CM272 treatment under both conditions. These results were also confirmed in mouse HCC cells NM53 and in human HCC cells **(Fig. 3H** and **Suppl. Fig. 3A, B)**.

**Figure 3.**
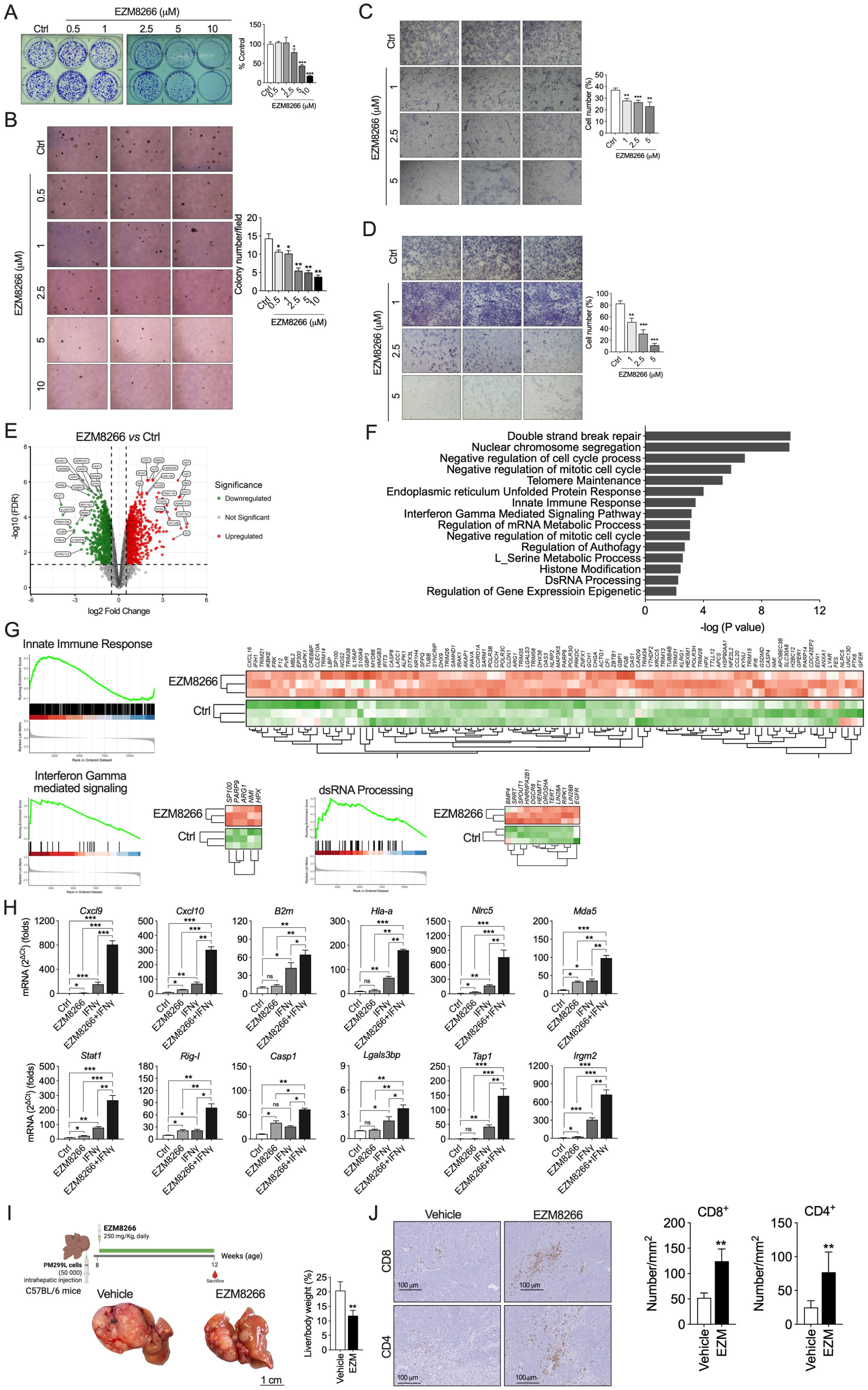
Antitumoral effects of the G9a-specific inhibitor EZM8266. (A) Colony formation assays performed in PLC/PRF/5 human HCC cells treated with the indicated concentrations of EZM8266. (B) Anchorage-independent growth assay performed in PLC/PRF5 cells treated with the indicated concentrations of EZM8266. Cell migration (C) and cell invasion (D) assays performed in HuH7 cells treated with the indicated concentrations of EZM8266 (48 h pretreatment and 24 h of transwell migration/invasion). (E) Volcano plot representing all the genes with significant differential expression (p<0.05) in PLC/PRF/5 HCC cells treated with EZM8266 (5 µM for 48 h) *vs* control cells, and most relevant categories of differentially expressed genes identified by Gene Ontology Biological Process (GO-BP) functional classification. (F) GSEA analysis of specific categories including heatmaps with a list of genes modulated by EZM8266 selected from the RNA-seq data. (G) Effect of EZM8266, INFγ and their combination on the expression of selected genes in the murine HCC cell line PM299L. (H) Experimental protocol for the study of the antitumoral effects of EZM8266 in an orthotopic model of HCC. Representative images of tumors and tumor burden (liver index) in vehicle and EZM8266-treated mice. (I) Representative images showing the immunohistochemical detection of CD4^+^ and CD8^+^ T cells and quantification of tumor-infiltrating CD8^+^ and CD4^+^ T cells (yellow arrows) in vehicle- and EZM8266-treated mice. * p<0.05, ** p<0.01, *** p<0.001.

Next, we examined the *in vivo* antitumoral properties of EZM8266 in an orthotopic liver tumor mouse model. PM299L cells were subcutaneously injected in mice and after 4 weeks tumors were retrieved, cut into 1-2 mm^3^ cubes and implanted into the left liver lobes of 2 groups of mice that were subsequently treated with EZM8266 or vehicle. We found that tumor growth was significantly reduced, and tumors’ weight at the end of treatments were also significantly lower **(Fig. 3I)**. Immunohistochemical analysis of CD8^+^ T cells tumoral infiltration revealed a higher level of this population in the smaller tumors found in mice treated with EZM8266 compared with those in control mice. Immunostaining for CD4^+^ T cells also revealed higher intratumoral levels following EZM8266 treatment compared with controls **(Fig. 3J)**.

### G9a inhibition suppresses immune evasion mechanisms in HCC cells by multiple mechanisms

According to our transcriptomic analyses in NM53 cells treated with CM272 and IFNγ, G9a inhibition seems to impinge on different mechanisms related to the activation of immunogenic pathways in HCC cells. In agreement with these findings, we observed that CM272 and IFNγ combination strongly enhanced CXCL10 release in both mouse and human HCC cells **(Fig. 4A** and **Suppl Fig. 4A)**. Furthermore, we also demonstrated that G9a inhibition led to a significant increase in cell surface expression of the MHC class I complex proteins. We validated these findings in cells treated with the G9a inhibitor EZM8266 **(Fig. 4B** and **Suppl Fig. 4B)**.

**Figure 4.**
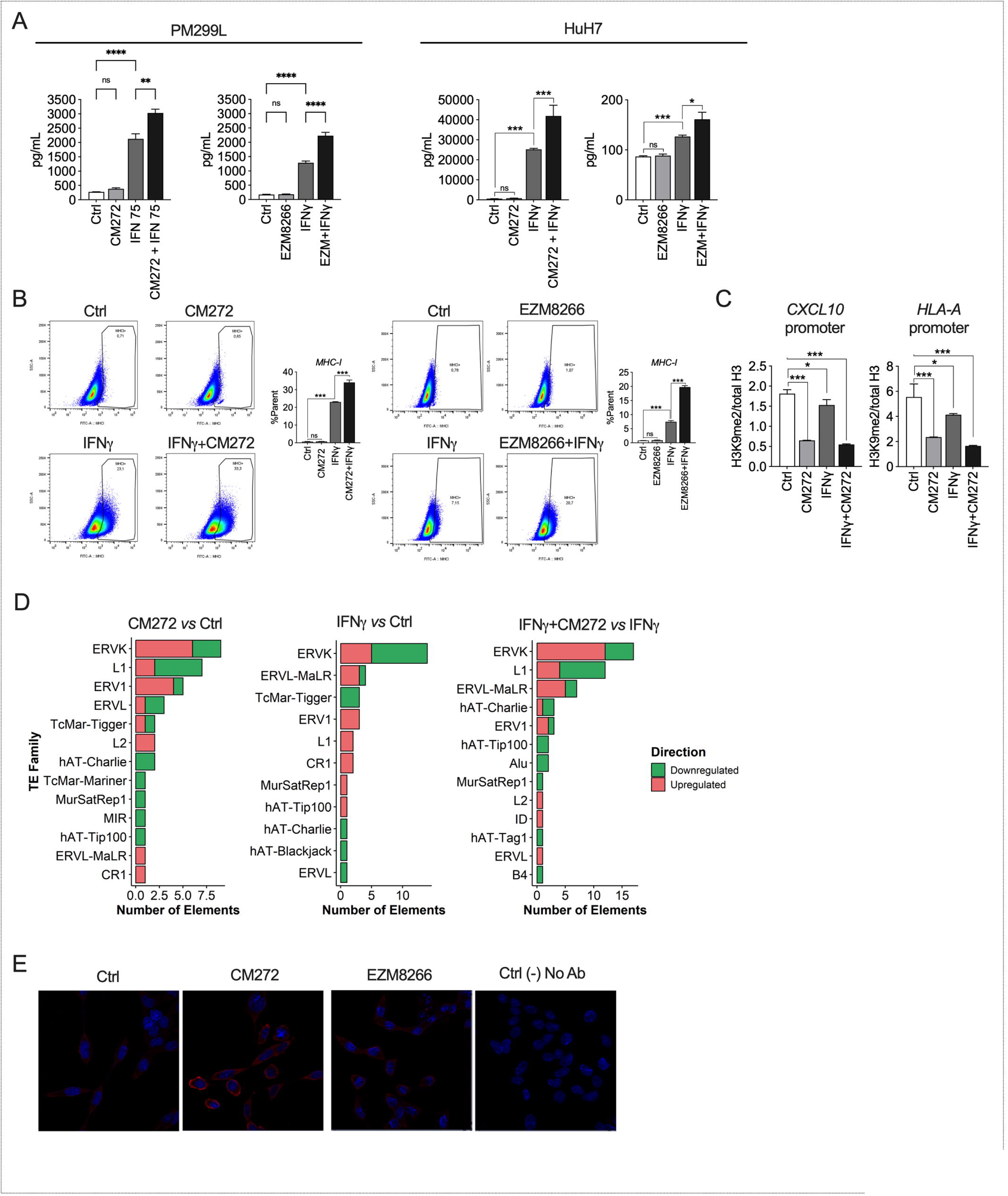
G9a inhibition potentiates the immunogenic effects of INFγ in HCC cells. (A) Effect of CM272 and EZM8266 on IFNγ-triggered CXCL10 production in murine (PM299L) and human (HuH7) HCC cells. Cells were treated with CM272 for 48 h or with CM272 for 24 h and then with INFγ (75 U/mL) for another 24 h, or with INFγ alone for 24 h. PM299L were treated with 400 nM and HuH7 received 1 µM of CM272. For EZM8266, cells were pretreated for 48 h with EZM8266 (5 µM) and then with IFNγ (75 U/mL) for another 24 h as indicated. CXCL10 protein levels were measured by ELISA in cells’ conditioned media. (B) Effect of G9a inhibition with CM272 or EZM8266 on the expression of MHC class I complex protein (MHC-I) on the surface of PM299L cells. Cells were treated with IFNγ and CM272 or EZM8266 as indicated in panel A, and MHC-I levels were determined by FACS analysis. (C) ChIP analyses of H3K9me2 levels in the proximal promoter regions of *CXCL10* and *HLA-A* genes in HuH7 cells treated with IFNγ (75 U/mL) and CM272 (400 nM) as indicated in panel A. (D) Evaluation of the expression of Transposable elements (TEs) and endogenous retroviral sequences (ERVs) by RNA-seq in NM53 murine HCC cells treated with IFNγ, CM272 and their combination as indicated in panel A. (E) Immunofluorescence analyses of dsRNA in PM299L HCC cells treated with CM272 (24 h) or EZM8266 for (48 h) h. Right panel shows a control without primary antibody. * p<0.05, ** p<0.01, *** p<0.001.

To gain more direct evidence on the role of G9a in the regulation of the expression of immune response-related genes, we performed ChIP analyses of the repressive G9a-mediated H3K9me2 mark in the promoter regions of relevant genes, such as *CXCL10* and the MHC class I gene *HLA-A*. As shown in **Fig. 4C**, the H3K9me2 mark, which was enriched in both *CXCL10* and *HLA-A* promoters, was significantly suppressed in HuH7 cells upon CM272 treatment. Moreover, CM272 enhanced the mild inhibitory effect that IFNγ has on this repressive epigenetic mark (**Fig. 4C**). These effects were reproduced in HuH7 cells treated with EZM8266 **(Suppl Fig. 4C)**.

Transposable elements (TEs) and endogenous retroviral sequences (ERVs) are DNA segments repressed through various epigenetic mechanisms, including those mediated by histone methyltransferases such as SETDB1^54^. When these epigenetic modifications are removed TEs and ERVs are transcribed, generating immunostimulatory double-stranded RNAs (dsRNAs)^55^. This response, termed viral mimicry, has been observed in the context of the inhibition of different epigenetic effectors, including DNA and histone-methyltransferases, in other epithelial cancers^56,57^. These host-derived dsRNAs mimicking viral dsRNA are detected by intracellular sensors such as RIGI and MDA5, triggering an interferon response^55^. Interestingly, RIGI was one of the most upregulated genes in HCC cells treated with G9a inhibitors plus IFNγ (**Fig. 2D**). We observed that the inhibition of G9a resulted in the upregulation of the expression of TE loci in NM53 mouse HCC cells, and that G9a targeting potentiated the response to INFγ. We observed induction of several endogenous retrovirus (ERVK, ERV1, ERVL) and LINE families, including L1, L2 or hAT transposable elements **(Fig. 4D)**.

To confirm that there is increased dsRNA formation in response to G9a inhibition, we performed immunofluorescence analyses of dsRNA in mouse and human HCC cells treated with G9a inhibitors in the presence or absence of IFNγ. We consistently observed that G9a inhibition, in cooperation with IFNγ stimulation, significantly increased the formation of dsRNA **(Fig. 4E** and **Suppl Fig. 4D)**. Taken together, these results indicate that specific G9a inhibition simultaneously triggers different immunostimulant mechanisms in HCC cells potentiating the response to IFNγ.

### G9a inhibition increases the efficacy of ICIs in clinically-relevant HCC models

In view of these compelling observations, we next investigated the effects of G9a inhibition on the therapeutic response to PD-1 blockade in an orthotopic mouse HCC model **(Fig. 5A)**. While EZM8266 monotherapy showed a significative, albeit limited effect, tumor growth was substantially abrogated when EZM8266 treatment was combined with anti-PD-1 antibodies **(Fig. 5B)**. Importantly, this therapeutic response was not accompanied by any observable side effects, including weight loss or internal organs abnormalities (data not shown). Moreover, the combination therapy significantly decreased the serum alanine transaminase, aspartate transaminase and lactate dehydrogenase levels to those found in age-matched normal mice **(Fig. 5C)**. Importantly, immunostaining analyses showed that EZM8266 or anti-PD-1 treatment induced CD8^+^ T cell infiltration in tumoral areas, and this response was significantly increased in mice treated with EZM8266 in combination with anti-PD-1 antibodies **(Fig. 5D)**. Similarly, an increase in CD4^+^ T cells was observed in the group of mice treated with the combination, although it did not reach statistical significance. These results suggest that G9a inhibition triggers an effective CD8^+^ T cell-mediated anti-tumor immunity, and provide a rationale for combining G9a inhibitors with ICIs to treat HCC.

**Figure 5.**
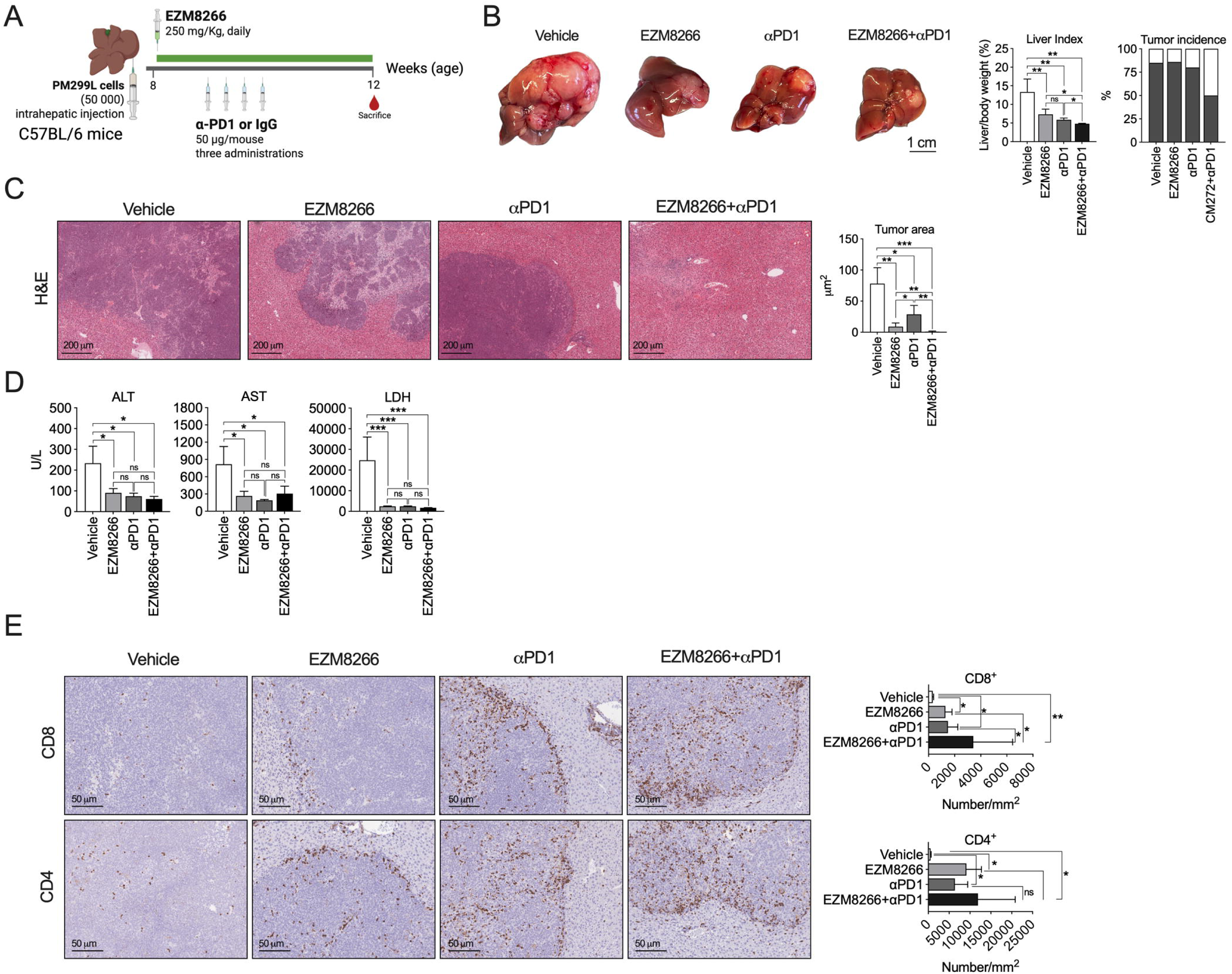
G9a inhibition with EZM8266 potentiates the in vivo antitumoral effect of ICI. (A) Experimental protocol for the study of the antitumoral effects of EZM8266 in combination with ICI in an HCC model developed by the orthotopic implantation of PM299L cells in immunocompetent mice. (B) Representative images of tumors in the different groups of mice at the end of treatments. Quantitation of tumor burden (liver index) and tumor incidence are indicated. (C) Histological evaluation of tumor growth (tumor area) in the different treatment groups. Representative images of H&E-stained liver and tumor tissues are shown. (D) Serum levels of ALT, AST and LDH in the different groups of mice at the end of treatments. (E) Representative images showing the immunohistochemical detection of CD4^+^ and CD8^+^ T cells and quantification of tumour-infiltrating CD8^+^ and CD4^+^ T cells in the different groups of mice at the end of treatments. * p<0.05, ** p<0.01, *** p<0.001.

### Effective combination of G9a inhibition and ICI in an aggressive recurrence-related HCC model

Approximately 30% of patients with HCC undergo resection or local ablation as primary treatment. However, the probability of tumor recurrence at 3 years is 30–50%^39^. Neoadjuvant and adjuvant administration of ICI have led to significantly improved outcomes compared to adjuvant treatment alone in patients with other tumor types, such as melanoma and non-small-cell lung cancer, which has prompted growing interest in exploring this strategy in HCC as well^4^. Surgical resection inherently causes tissue injury, triggering processes of wound healing and inflammation^58^. The regenerative response following a partial hepatectomy (PH) involves a complex interplay of growth factors, reactive oxygen species, and pro-inflammatory cytokines. These mediators are released during and shortly after surgery and are hypothesized to contribute to tumor recurrence by modulating the tumor microenvironment^59,60^. Building on the favorable outcomes observed in our previous experimental models, we aimed to investigate whether G9a inhibition could enhance the efficacy of ICI therapy in the context of post-hepatectomy inflammation. To this end, we employed an orthotopic model of HCC combined with PH, designed to evaluate tumor progression in a regenerative liver milieu. In this model, HCC cells were injected into one of the remaining liver lobes following resection **(Fig. 6A)**. Monotherapy with CM272 induced a moderate antitumor effect, while anti-PD-1 antibody administration led to a noticeable reduction in tumor burden. Strikingly, the combination of CM272 and anti-PD-1 therapy resulted in marked tumor regression **(Fig. 6B, C)**. In alignment with our previous findings using EZM8266, this combinatorial approach also reshaped the immune cell landscape within tumor tissues. Specifically, the proportion of tumor-infiltrating CD8⁺ T cells was significantly increased in CM272-treated mice compared to controls, with further enhancement observed upon anti-PD-1 co-treatment. Likewise, CD4⁺ T cell infiltration was significantly elevated in the combination treatment group relative to controls (**Fig. 6E**). Taken together, these results confirmed for the first time in HCC how G9a inhibition can enhance the sensitivity of anti-PD-1 therapy inducing CD8^+^ T cell-mediated antitumor immunity, constituting a promising combinatorial strategy against this tumor in different scenarios.

**Figure 6.**
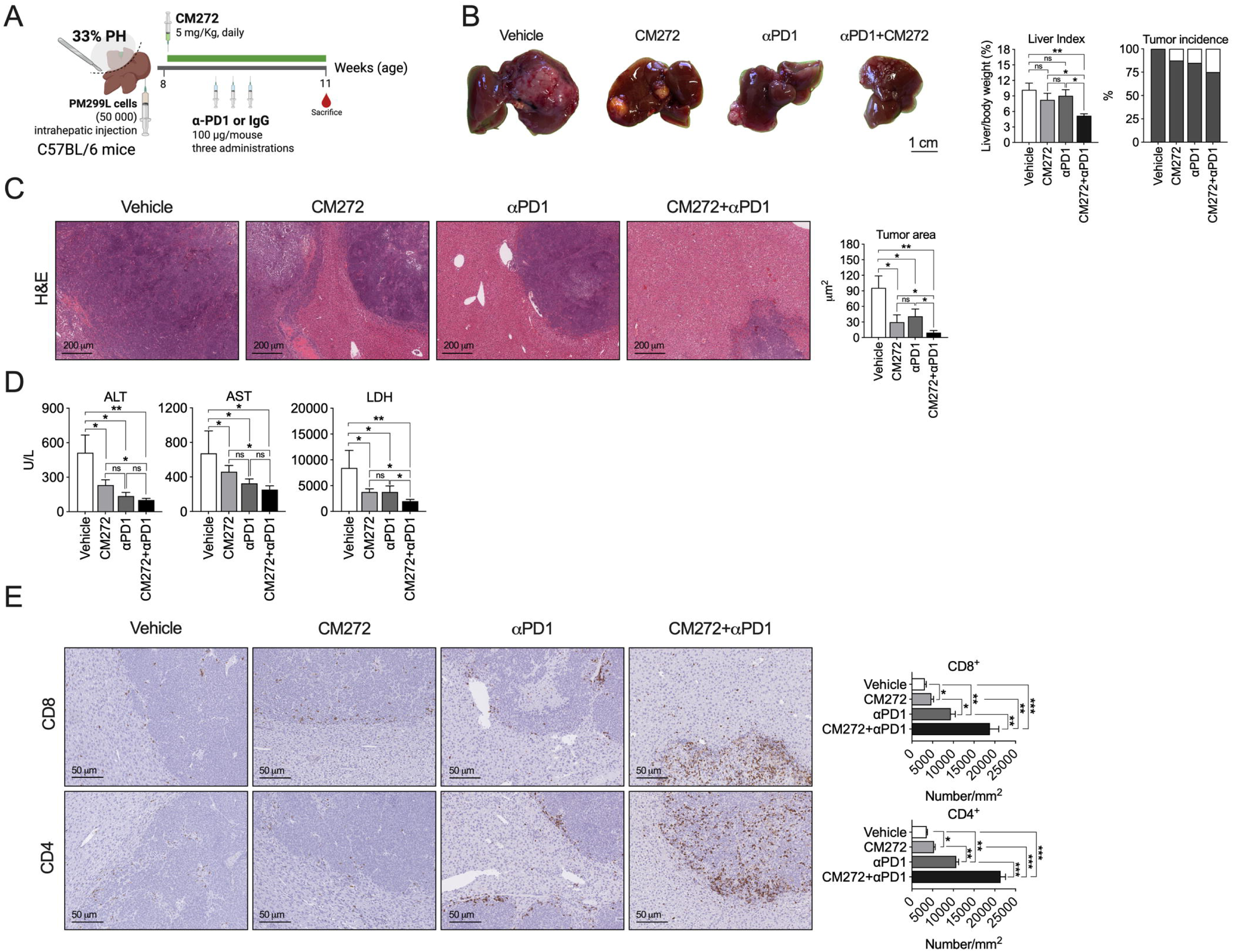
G9a inhibition with CM272 potentiates the in vivo antitumoral effect of ICI in an aggressive model of PH-stimulated HCC growth. (A) Experimental protocol for the study of the antitumoral effects of CM272 in combination with ICI in a HCC model developed by the orthotopic implantation of PM299L cells in immunocompetent mice, and a concomitant partial hepatectomy (33% PH). (B) Representative images of tumors in the different groups of mice at the end of treatments. Quantitation of tumor burden (liver index) and tumor incidence are indicated. (C) Histological evaluation of tumor growth (tumor area) in the different treatment groups. Representative images of H&E-stained liver and tumor tissues are shown. (D) Serum levels of ALT, AST and LDH in the different groups of mice at the end of treatments. (E) Representative images showing the immunohistochemical detection of CD4^+^ and CD8^+^ T cells and quantification of tumor-infiltrating CD8^+^ and CD4^+^ T cells in the different groups of mice at the end of treatments. * p<0.05, ** p<0.01, *** p<0.001.

## Discussion

Immunotherapy has revolutionized cancer treatment in the past decade, with ICI becoming a cornerstone of therapy in various malignancies^61^. However, in HCC, the clinical benefit of ICI remains limited to a subset of patients^3^. This underscores the pressing need to identify novel molecular targets and rational combination strategies to enhance immunotherapeutic efficacy in this setting. The inherent ability of HCC to escape immune surveillance presents a major challenge for the successful implementation of immunotherapy in clinical practice^4^. In this context, tumor cells exploit both genetic and epigenetic mechanisms to evade immune detection and destruction. Recent studies have highlighted the role of epigenetic regulators in modulating antitumor immunity, positioning them as attractive combinatorial partners for ICI. Although relatively few studies have investigated this in HCC specifically, inhibitors targeting DNA methyltransferases (DNMTs), histone deacetylases (HDACs), the histone methyltransferase EZH2, or bromodomain and extraterminal domain (BET) proteins have demonstrated the potential to enhance ICI efficacy^14^. Nevertheless, a comprehensive preclinical validation remains necessary before these strategies can be clinically implemented. In the present study, we provide evidence that the histone methyltransferase G9a can play a significant role in modulating resistance to immunotherapy in HCC. Using an integrative approach that included the TCGA cohort and four independent HCC transcriptomic datasets, encompassing a total of 764 patients, we observed that high G9a expression inversely correlated with relevant gene expression signatures predictive of ICI response. We first assessed the relationship between G9a expression and established HCC molecular subclasses, including the Inflamed class and the IFN18 signature. In addition, we evaluated its association with a series of immune gene signatures known to distinguish responders from non-responders to ICI, such as IFNAP, ABRS, and more recently developed single-cell-derived signatures including CD8Tex, CD8Temra, and Macro-CXCL10^45^. Strikingly, patients with low scores across these immune signatures consistently displayed elevated G9a expression. These findings suggest that G9a overexpression may contribute to an immunosuppressive tumor microenvironment and impaired response to ICI in HCC. Thus, targeting G9a could represent a novel strategy to restore immune surveillance and overcome resistance to immunotherapy in this neoplasia. Importantly, the role of G9a in modulating immune responses has been previously explored in preclinical models of several tumor types, including melanoma^53^, ovarian cancer^62^, glioma^63^, head and neck squamous cell carcinoma^64^, and bladder cancer^65^ with promising results. Our study expands this growing body of evidence by demonstrating its clinical relevance in patients with HCC. Interestingly, the role of G9a has been previously examined in the immune compartment as an epigenetic regulator influencing T cell cytotoxicity^66^. In a study utilizing engineered T cells targeting HCC, short-term inhibition of G9a enhanced T cell antitumor activity both *in vitro* and in an orthotopic mouse model. Specifically, G9a blockade increased granzyme expression without promoting terminal T cell differentiation or exhaustion, and induced selective transcriptional and proteomic changes in genes involved in pro-inflammatory signaling, T cell activation, and cytotoxicity^66^. These findings are highly consistent with our observation that G9a expression in HCC patients inversely correlates with a wide range of immune-related genes implicated in these mentioned processes. These include those involved in cytolytic activity (e.g., *GZMA, GZMB, GZMK, GZMH, PRF1*), inflammation-initiation chemokines (*CXCL9, CXCL10, CXCL11, CCL2, CCL4, CCL5*), T cell markers (*CD3D, CD3E, CD3G, CD8A*), NK cell cytotoxicity (*NKG7, KLRD1, KLRK1*), antigen presentation (*CIITA, HLA-DRA, B2M, PTPRC*), as well as other immunomodulatory molecules such as *IFITM1, PRKCH*, and *GBP1*. Collectively, these data provide a compelling rationale for exploring the specific role of G9a in HCC tumor cells and for the development of innovative G9a-targeting agents as part of combination immunotherapy strategies in HCC. Currently, a number of small-molecule inhibitors have been developed to target G9a with acceptable, though limited, therapeutic efficacy. However, most have not progressed to clinical trials due to suboptimal physicochemical and pharmacokinetic properties^67^. In the present work, we investigated the molecular consequences of G9a inhibition in murine HCC cells using two novel and promising inhibitors: CM272, a selective, reversible chemical probe with previously demonstrated antitumor activity in HCC models^16^, and EZM8266, a highly selective G9a inhibitor in advanced stages of preclinical development^21^. Our data demonstrate–that EZM8266 exerts potent antitumor effects in HCC. Functional assays—including colony formation, anchorage-independent growth, and transwell migration/invasion—revealed that EZM8266 treatment significantly impaired HCC cell clonogenicity, migration, and invasiveness in vitro. Moreover, orthotopic tumor models confirmed that *in vivo* administration of EZM8266 substantially reduced tumor growth in immunocompetent mice, with no signs of systemic toxicity and an excellent safety profile. Mechanistically, our data demonstrate that G9a inhibition in HCC tumor cells disrupts multiple layers of immune evasion. Pharmacologic targeting of G9a using two structurally distinct inhibitors, CM272 and EZM8266, induced broad transcriptional reprogramming characterized by the activation of interferon signaling and other immune-associated pathways. In both murine and human HCC cells, G9a inhibition synergized with IFN-γ to enhance the transcription of immune effector genes, including those encoding chemokines such as CXCL10 and components of the major histocompatibility complex class I (MHC-I), thereby promoting both lymphocyte recruitment and tumor cell visibility to cytotoxic immune cells. ELISA assays confirmed a robust increase in IFN-γ–induced CXCL10 secretion upon treatment with either CM272 or EZM8266. In parallel, our flow cytometric analyses revealed significantly enhanced surface expression of MHC-I molecules on HCC cells, particularly under IFN-γ stimulation, suggesting that G9a inhibition facilitates antigen presentation. Chromatin immunoprecipitation assays demonstrated a marked reduction of the repressive histone mark H3K9me2 at the promoters of CXCL10 and HLA-A, indicating that G9a inhibition epigenetically primes chromatin for transcriptional activation. In addition to reprogramming immune-related gene expression, G9a inhibition consistently upregulated transcripts derived from transposable elements (TEs), particularly endogenous retroviruses (ERVs). This observation is in line with prior studies demonstrating that inhibition of epigenetic repressors such as DNMT1^68^, SETDB1^69,70^, or even G9a in other types of tumors ^21,68^, can derepress ERVs, leading to their transcriptional activation. These sequences, remnants of ancient viral integrations into the host genome, can trigger innate antiviral responses via the induction of double-stranded RNA (dsRNA) and the subsequent activation of viral defense pathways—a phenomenon termed “viral mimicry”^55^. In our study, G9a inhibition induced a significant accumulation of immunostimulatory dsRNA in both mouse and human HCC cells, further supporting the engagement of viral mimicry as a key mechanism for enhancing tumor immunogenicity.

Collectively, these data establish that G9a inhibition not only reverses epigenetic silencing of immune-stimulatory genes and TEs, but also fundamentally increases tumor immunogenicity. This mechanistic insight provided the rationale for combining G9a inhibitors with immune checkpoint blockade. Indeed, combinatorial treatment with either CM272 or EZM8266 and anti–PD-1 antibodies led to near-complete regression of orthotopically implanted HCC tumors, including in aggressive post-hepatectomy models, without signs of hepatic or systemic toxicities. This effect was accompanied by a significant increase in intratumoral CD8⁺ T cell infiltration, reinforcing the role of G9a inhibition in enhancing cytotoxic T cell–mediated tumor clearance. While our mechanistic investigations centered primarily on tumor-intrinsic epigenetic alterations and their effects on CD8⁺ T cells, the broader impact of G9a inhibition on other immune subsets remains to be elucidated. Future studies should explore the effects on natural killer (NK) cells, dendritic cells, myeloid-derived suppressor cells (MDSCs), and regulatory T cells (Tregs), which may also play critical roles in shaping antitumor immunity.

In summary, our findings implicate G9a as a central regulator of immune escape in HCC and a contributor to resistance to immune checkpoint inhibition. G9a inhibition not only enhances the immunogenicity of HCC cells through transcriptional and epigenetic remodeling but also synergizes with IFN-γ to amplify antigen presentation and chemokine-driven immune cell recruitment. When combined with anti–PD-1 therapy, G9a inhibition results in potent antitumor activity and durable immune responses *in vivo.* Together with previous reports demonstrating enhanced CD8⁺ T cell cytotoxicity upon G9a inhibition, our study supports a compelling rationale for integrating G9a-targeting agents into combination immunotherapy strategies. Given the efficacy, potency, and favorable safety profile of CM272 and EZM8266, these agents represent promising candidates for clinical development as adjuncts to current immunotherapeutic regimens in HCC^69^

---

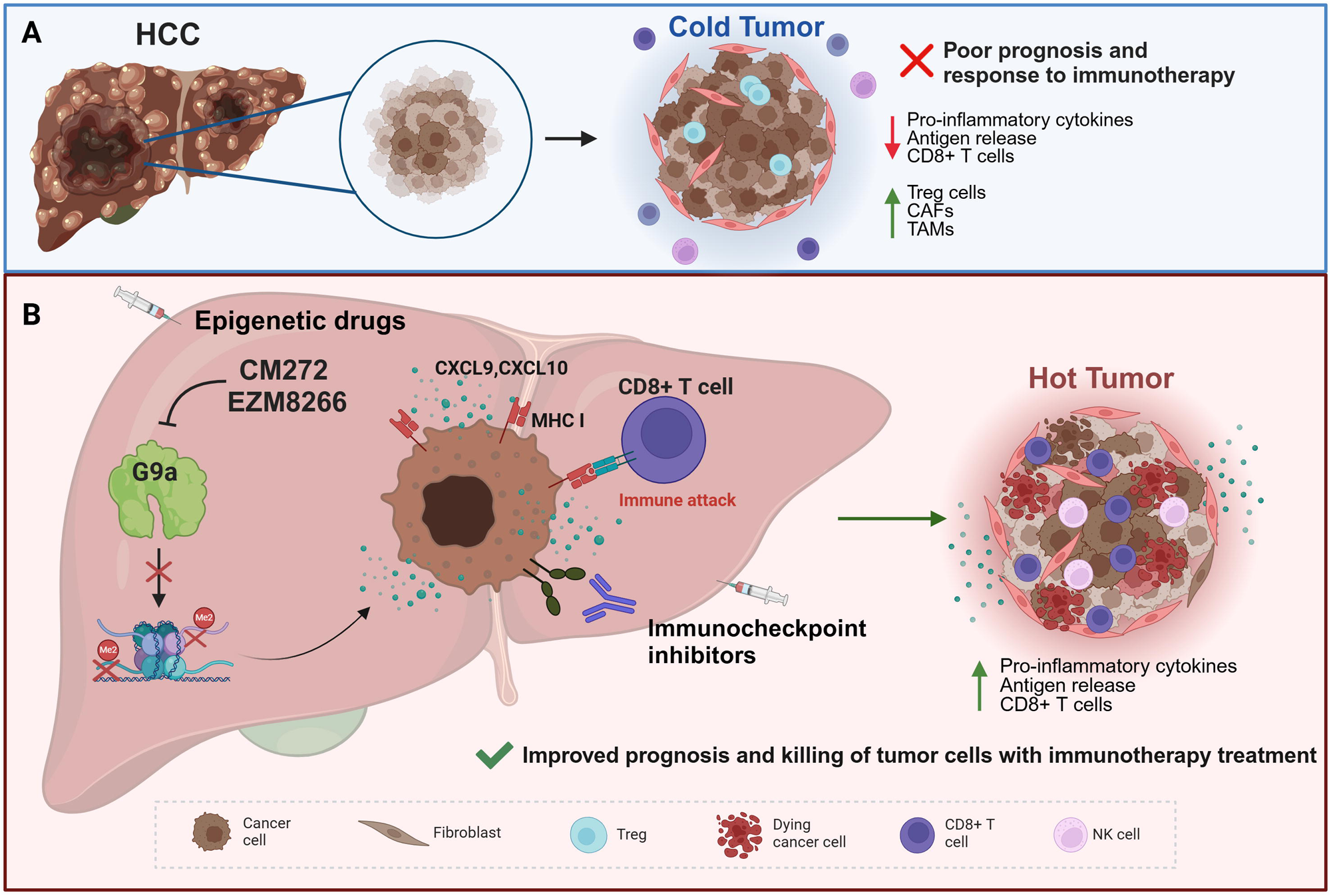

