## Supplementary figures and images for "Histone methyl-transferase G9a inhibition boosts the efficacy of immune checkpoint inhibitors in experimental hepatocellular carcinoma"

### Suppl Fig. 1A

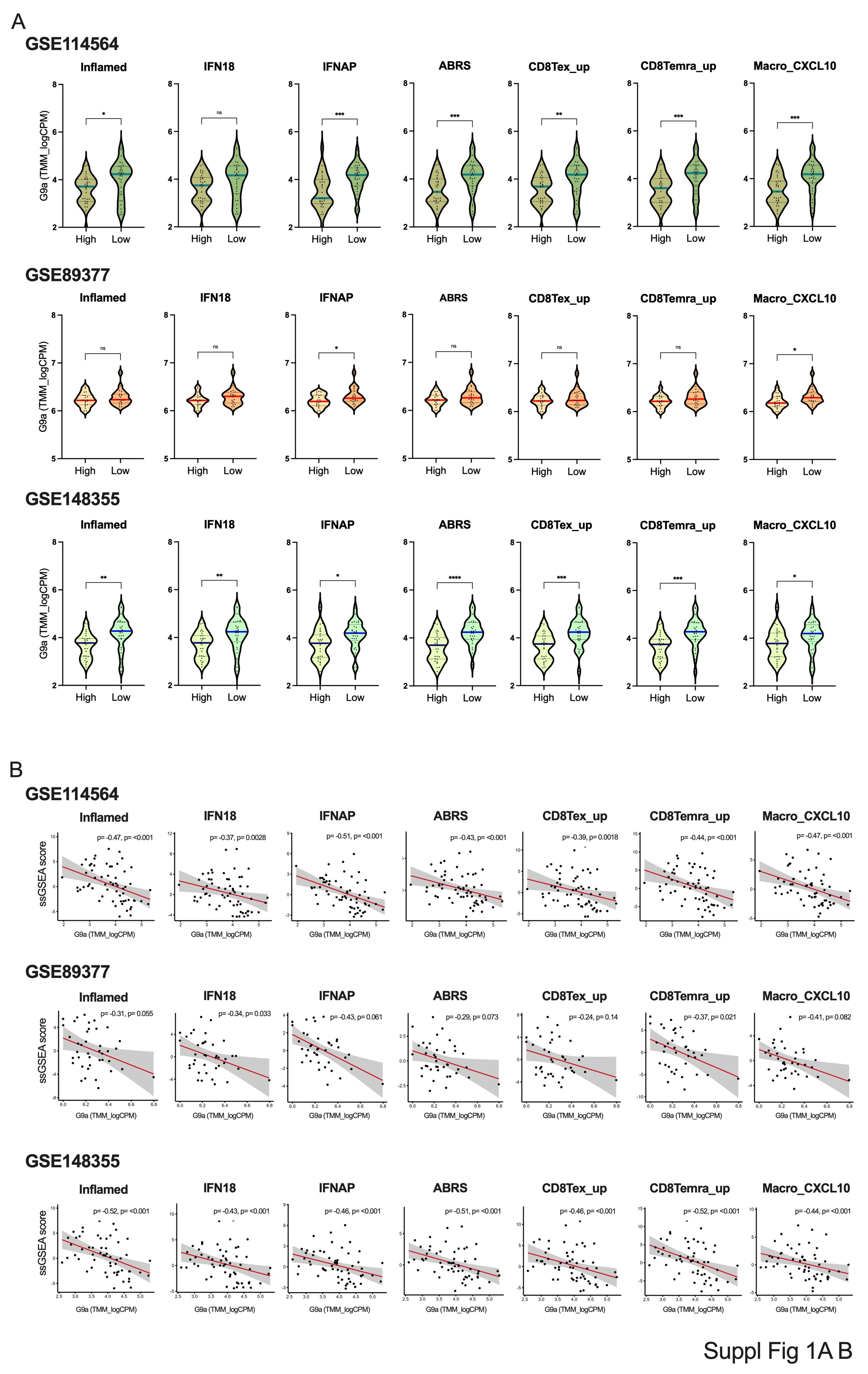

### Suppl Fig. 1C

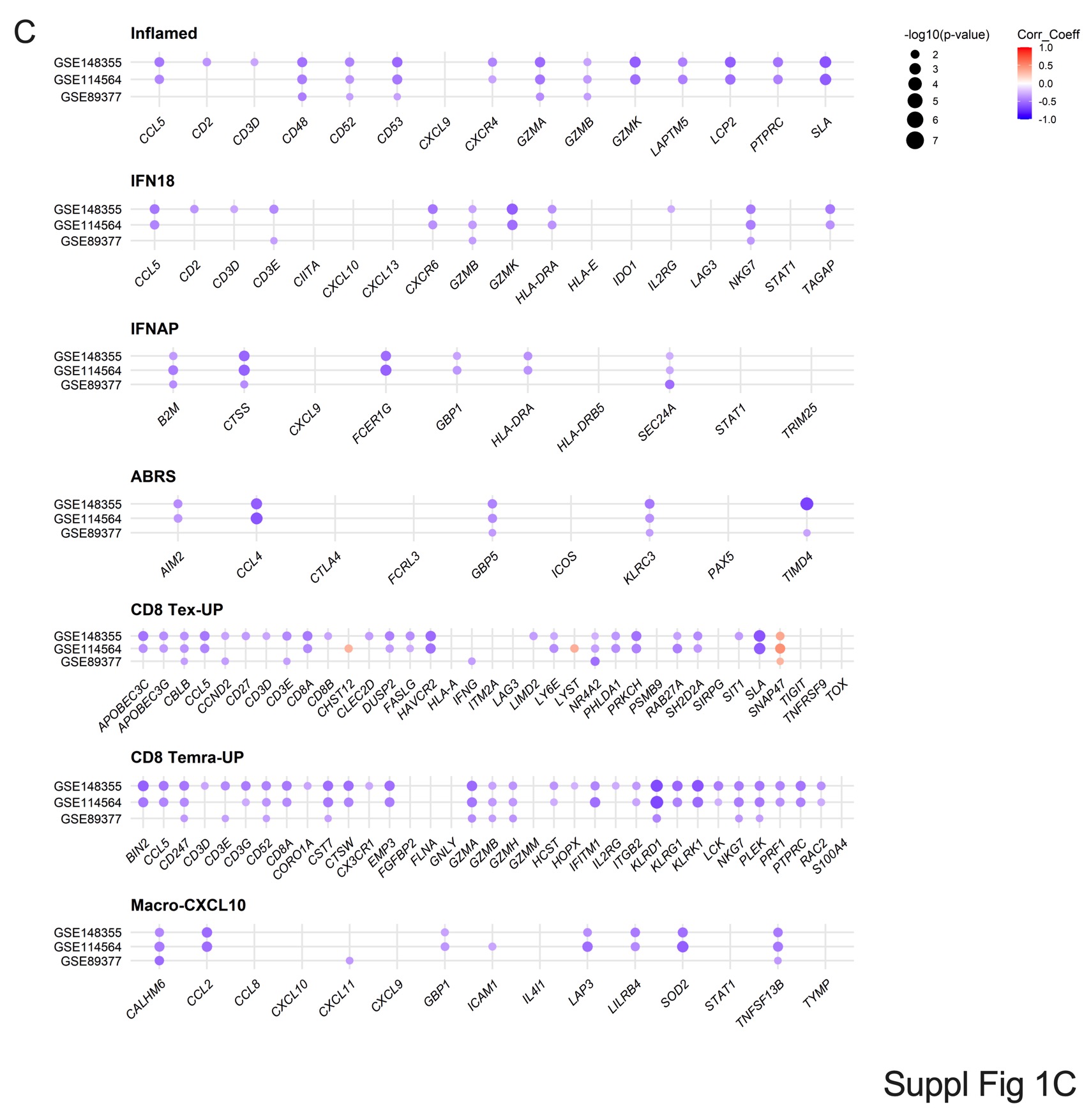

### Suppl Fig. 2

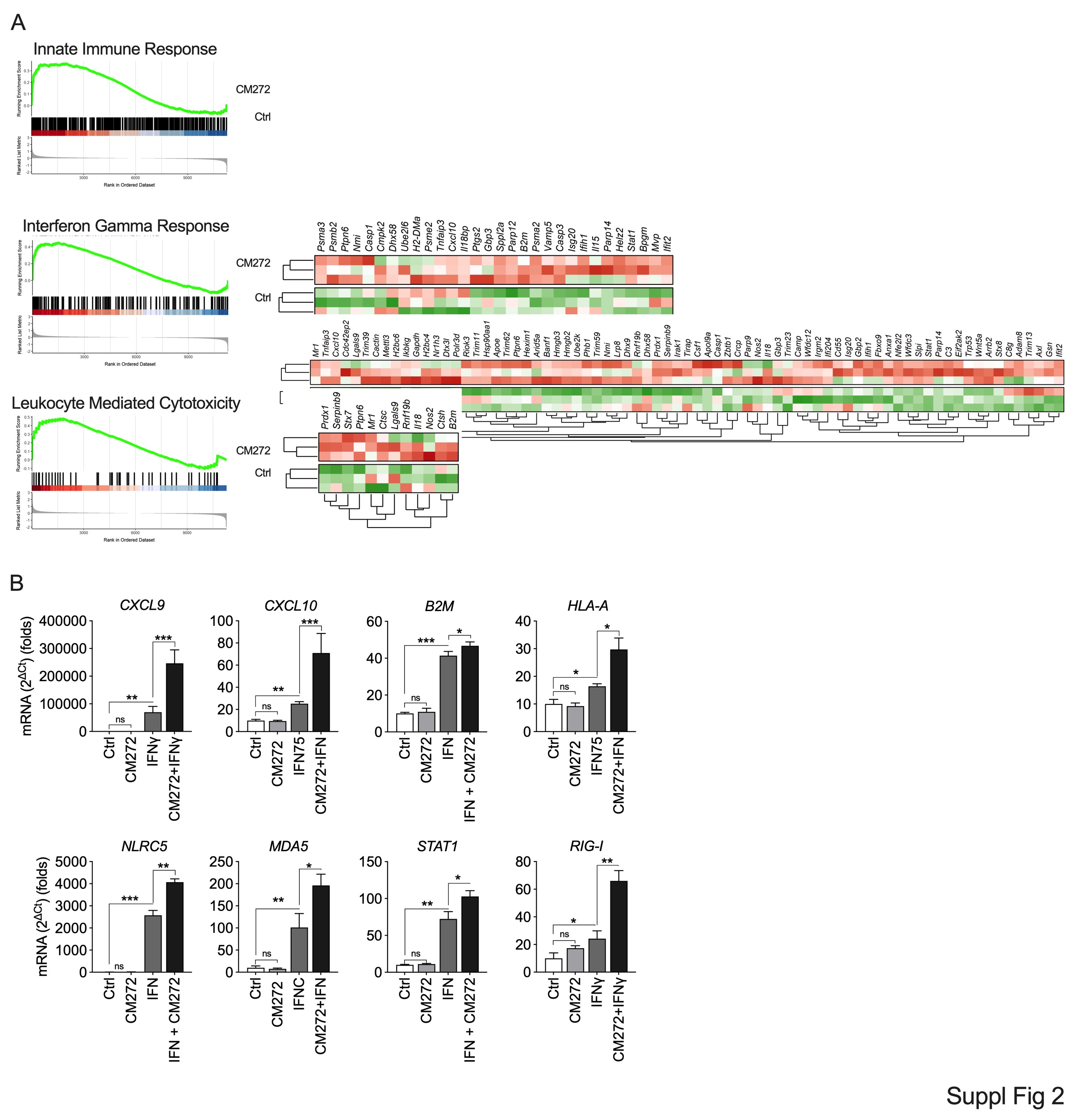

### Suppl Fig. 3

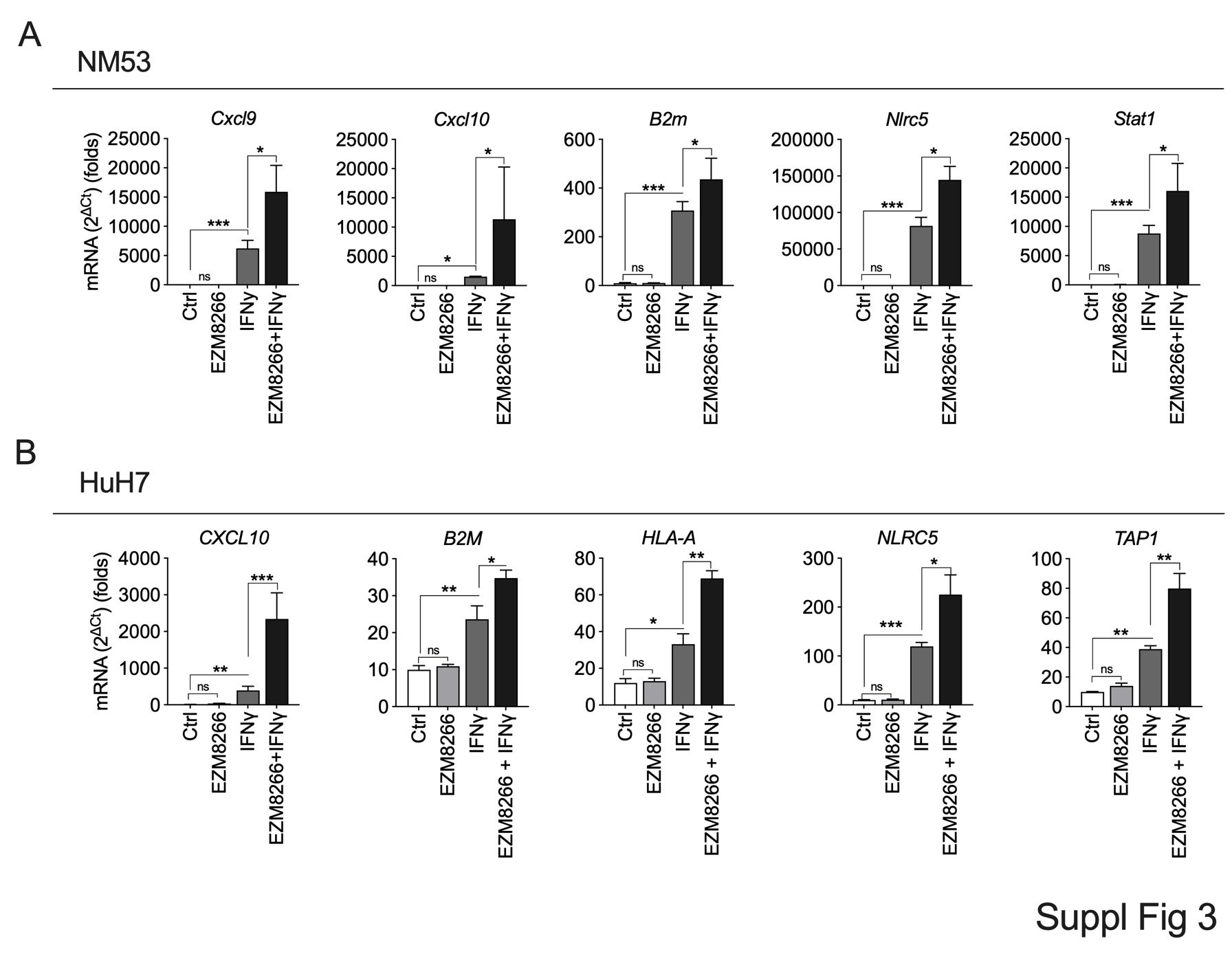

### Suppl Fig. 4

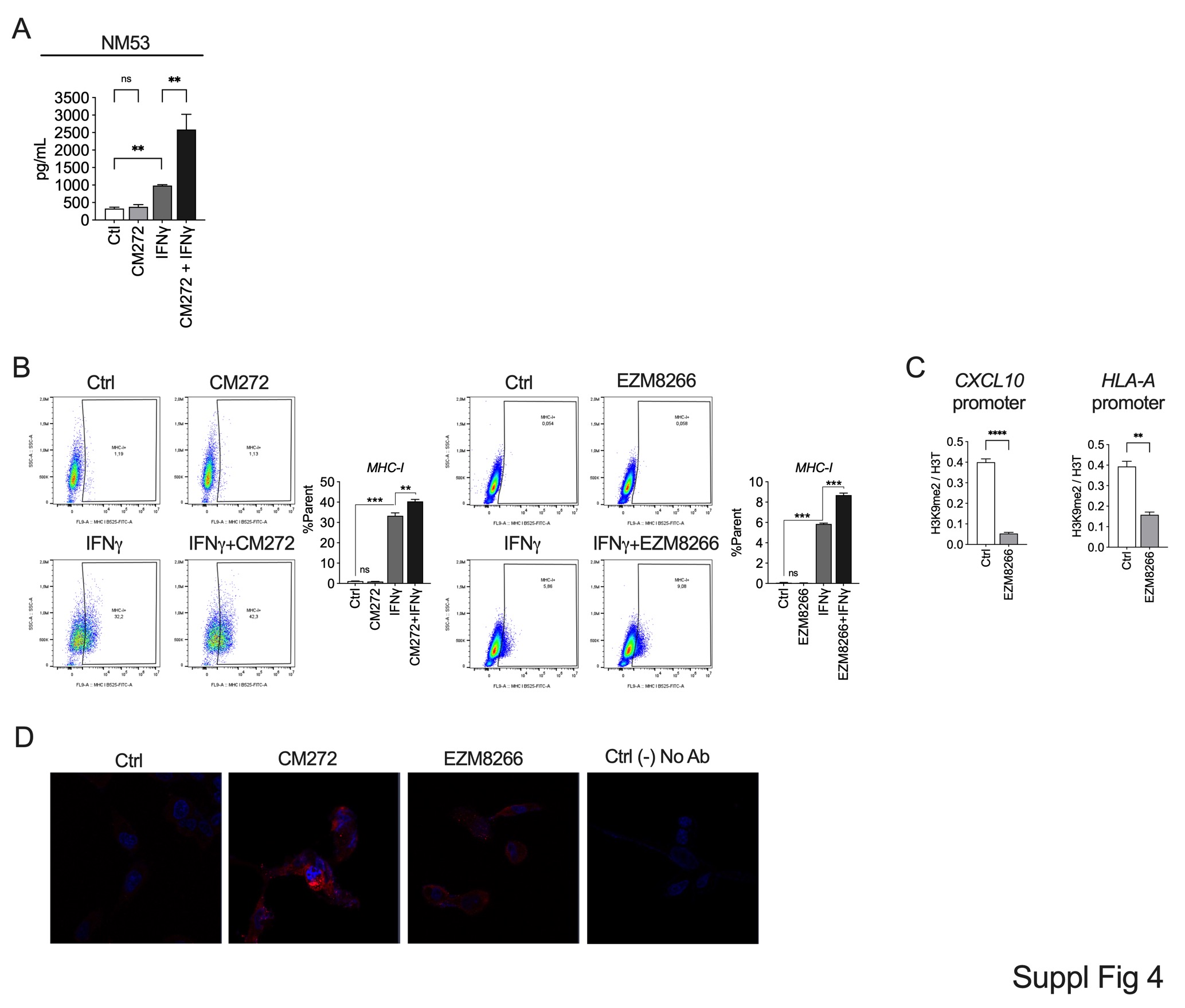
